# Regional adaptation to mosquito vectors shapes *Plasmodium falciparum* populations

**DOI:** 10.1101/2025.11.13.688255

**Authors:** Duangkamon Loesbanluechai, Lauriane Sollelis, Megan Armstrong, Inish Menezes, Amelia Cox, Lizzie Bridget Tchongwe Divala, Anne Lawson, Sabyasachi Pradhan, Prince Chigozirim Ubiaru, Antrea Pallikara, Dorothy Armstrong, Gillian Parker, Chiyun Lee, Richard D Pearson, Barbara H Stokes, Lisa Ranford-Cartwright, Matthias Marti, Virginia Howick

## Abstract

Transmission of *Plasmodium falciparum* through mosquitoes represents the most severe population bottleneck in the parasite’s life cycle, yet the genetic basis of parasite–vector compatibility remains poorly understood. Here, we show that mosquito species–specific transmissibility depends on allelic variation in multiple *P. falciparum* genes expressed during midgut invasion, beyond the well-studied Pfs47. Using an allelic replacement strategy, we targeted highly geographically differentiated SNPs in *P. falciparum* that match regional variation in vector community composition. Transmissibility was compared across four mosquito species representing distinct geographic ranges (*An. gambiae*, *An. stephensi*, *An. minimus*, and *An. albimanus*). Two of five tested polymorphisms showed increased oocyst and sporozoite burdens in sympatric parasite–vector combinations compared to allopatric ones. Both substitutions occurred in ookinete micronemal proteins, CTRP and WARP, within von Willebrand factor A domains, suggesting that regional allelic variation modulates *Plasmodium*–vector compatibility by altering midgut adhesion interactions. These findings reveal that vector compatibility is a polygenic trait shaped by molecular interactions across several loci. Understanding this complexity refines models of parasite adaptation and can inform the design of transmission-blocking interventions effective across diverse vector-parasite combinations.

## Introduction

Malaria remains one of the world’s most devastating infectious diseases, causing more than 500,000 deaths each year [1]. The disease is caused by *Plasmodium* parasites transmitted through the bites of *Anopheles* mosquitoes. Among these parasites, *Plasmodium falciparum* is the most virulent and is widespread across the tropics. However, the community composition of *Anopheles* vectors varies geographically and may impose strong, regionalized selective pressures on the parasite [2–4].

Transmission through the mosquito represents the most severe population bottleneck in the *Plasmodium* life cycle, making it a critical target for transmission blocking interventions. After being ingested during an infectious blood meal, parasites must undergo rapid sexual differentiation, fertilization, and meiosis before forming invasive ookinetes that traverse the mosquito midgut epithelium [5]. Only a small fraction of ookinetes successfully cross this barrier: parasite numbers often drop from thousands of gametocytes to fewer than ten oocysts [6]. In incompatible vectors, parasites are eliminated by the mosquito immune system or fail to exit the midgut lumen [7,8].

A key molecular determinant of this compatibility is the parasite antigen Pfs47, which mediates evasion of the mosquito immune response. Variants of Pfs47 act as the “key” that must match a mosquito “lock” for successful infection [9]. Population genomic analyses reveal strong geographic structuring, with near fixation of distinct Pfs47 haplotypes across major continental regions, consistent with adaptation to regional vector communities [9–12]. However, genome-wide analyses indicate that many other *P. falciparum* genes expressed during mosquito transmission show similarly strong geographic differentiation [13,14]. These patterns suggest that vector compatibility may be a polygenic trait, shaped by multiple interacting loci rather than a single lock-and-key mechanism.

Here, we tested this hypothesis by generating allelic replacement lines for five geographically differentiated transmission-stage genes and assaying their infectivity across four *Anopheles* species with distinct geographic ranges. We found that for two genes, sympatric parasite–vector combinations achieved higher transmission success than allopatric pairings. Both loci encode micronemal proteins, with substitutions localized to the same adhesive functional domain, suggesting direct interactions with mosquito midgut components. These findings support a model of vector compatibility driven by coevolution between *Plasmodium* and local mosquito populations through multiple molecular interfaces.

## Results

### Identification and allelic replacement of candidate loci

To identify loci involved in adaptation to regional mosquito vectors, we integrated population genomic and gene expression data to pinpoint variants that may influence parasite infectivity in the mosquito midgut. The MalariaGEN network analysed genomic variation in 7,000 *P. falciparum* samples and ranked genes by a global differentiation score based on the most divergent non-synonymous SNP per gene [13]. We selected the top 5% of genes (278) from this list and cross-referenced them with Malaria Cell Atlas expression data [15–17] to identify those associated with gametocytes- or ookinete- specific gene clusters and/or were upregulated in *P. falciparum* relative to *P. berghei* during the ookinete stage (844 genes). These expression criteria enriched for genes likely involved in midgut traversal, and we hypothesized that transcriptional differences between *P. falciparum* and *P. berghei* reflect vector-specific adaptations to their respective mosquito hosts [18]. This intersection yielded 60 candidates (**Table S1**), from which eight were selected (**Table 1**), on the basis of high expression in mature gametocytes or ookinetes, and presence of high-frequency alternative alleles in geographic regions that coincided with mosquito species available in our insectary (**Table 1, S1; Figure 1A, S1, S2**). Six of the genes (CTRP, Pfs25, Pfs47, SOAP, WARP, P230) have established roles in transmission and are targets for potential transmission-blocking interventions [19–22]. P230, P25, and P47 are surface proteins involved in gamete interaction, midgut traversal, and immune evasion [11,23–26]. SOAP, WARP, and CTRP localize to ookinete micronemes and mediate motility and midgut invasion [27–31], with CTRP knockout blocking parasite invasion of the mosquito midgut [32]. The remaining two loci, HSP101 and Pf3D7_1004600, have not previously been linked to transmission but show higher expression in *P. falciparum* ookinetes compared to *P. berghei* [15]. This species-specific expression suggests possible adaptation to distinct mosquito hosts. HSP101 encodes a PTEX translocon component active in asexual blood stages [33], while Pf3D7_1004600 encodes a protein of unknown function.

**Figure 1.**
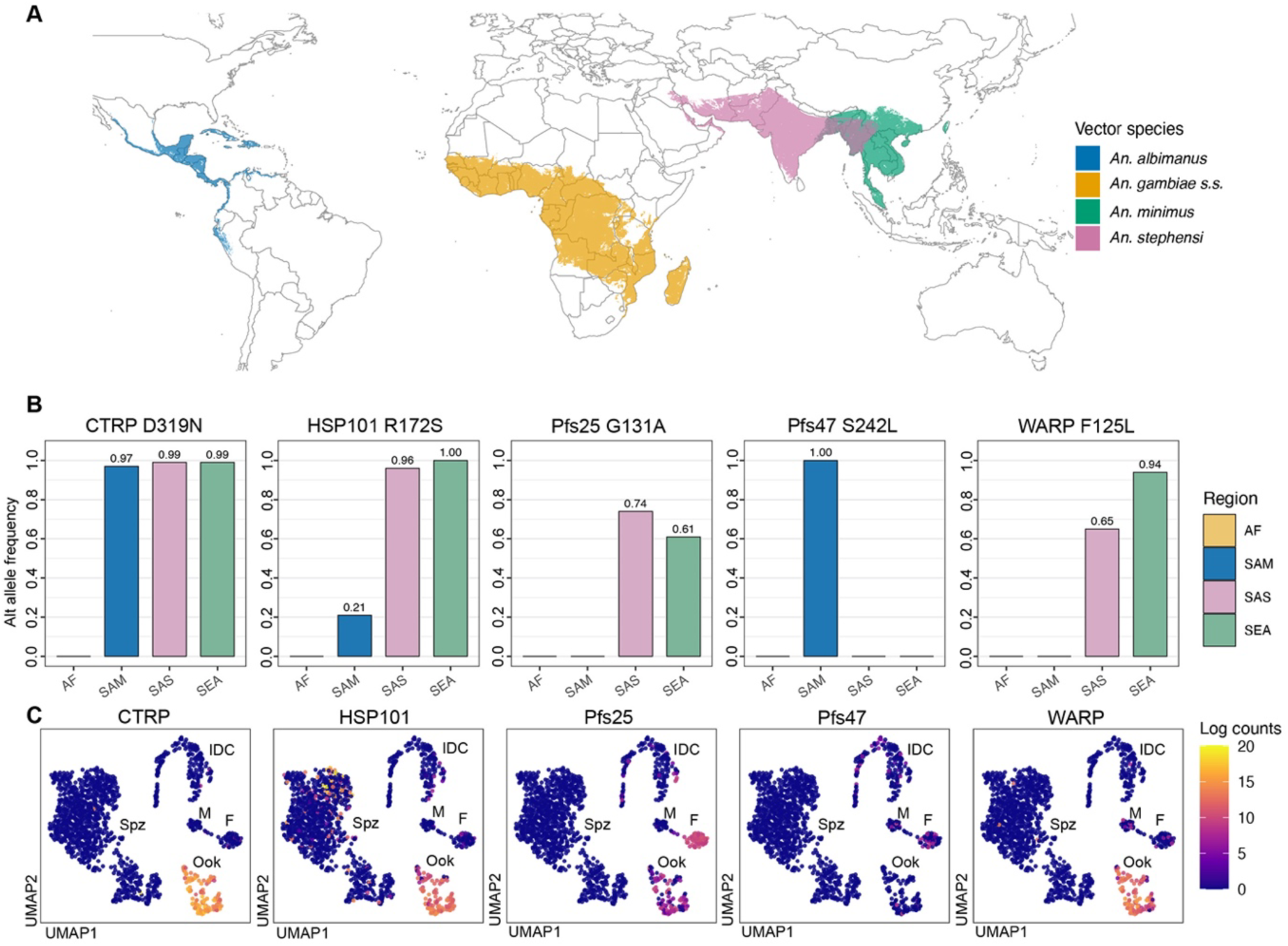
Distribution of *Anopheles* species and *P. falciparum* genotypes. (A) The global distributions of *An. albimanus, An. gambiae, An. minimus,* and *An. stephensi* from the Malaria Atlas Project [2,34]. Shaded areas indicate the probability of presence is greater than 50%. (B) Alternative allele frequencies for the five candidate loci across four broad geographic areas associated with each of the vector species: Africa (AF), South America (SAM), South Asia (SAS), Southeast Asia (SEA). Data are from [35]. We defined a sympatric combination of parasite allele and vector species as a co-occurrence of the alternative allele at greater than 50% frequency in the broad geographic area and presence of the vector in at least one country within that region where parasite genomic information was available (See also **Figure S2**). (C) UMAP of single-cell gene expression data from the Malaria Cell Atlas project [15] for the five candidate genes. High expression was observed in ookinetes and/or gametocytes for each gene (IDC = Intraerythrocytic Developmental Cycle; M = Male gametocyte; F = Female gametocyte; Ook = Ookinete; Spz = Sporozoite).

**Table 1.** List of candidate loci. Eight candidate genes were identified for allelic replacement. Shown are the gene ID, gene name, amino acid (AA) change, RNA expression pattern of the gene, global differentiation score with rank, and whether transfection was successful (**✓**) or not (✗). Gene expression data were analysed from the Malaria Cell Atlas [15,16] (**Figure S1**). Global differentiation score, a gene-level *Fst*-derived metric, and rank out of the 5,561 *P. falciparum* genes are from the MalariaGEN Pf6 dataset [13].

| Gene ID | Gene name | AA change | Highest gene expression stage | Global differentiation score (rank) | Transfection (3D7) |
| --- | --- | --- | --- | --- | --- |
| PF3D7_1346800 | Pfs47 | S242L | Gametocyte | 1 (1) | ✓ |
| PF3D7_0315200 | CTRP | D319N | Ookinete | 0.98 (5) | ✓ |
| PF3D7_1404300 | SOAP | S95R | Ookinete | 0.62 (120) | ✗ |
| PF3D7_0801300 | WARP | F125L | Ookinete | 0.66 (64) | ✓ |
| PF3D7_0209000 | P230 | N1566T | Gametocyte | 0.61 (141) | ✗ |
| PF3D7_1031000 | Pfs25 | G131A | Ookinete | 0.58 (205) | ✓ |
| PF3D7_1116800 | HSP101 | R172S | Ookinete | 0.81 (7) | ✓ |
| PF3D7_1004600 | . | L94I | Ookinete | 0.67 (53) | ✗ |

To assess the role of these non-synonymous SNPs in mosquito-specific adaptation, we used CRISPR-Cas9 to generate transgenic *P. falciparum* parasites by inserting the alternative allele into the reference parasite strain 3D7, which is of African origin, to generate a single amino acid change in each gene (**Figure S3**). For each gene, we also generated a synonymous (silent) control line that recoded the gRNA/PAM region to prevent Cas9 re-cutting while leaving the protein sequence unchanged, serving as a transgenesis-handling control for our phenotypic assays (**Table S2**). Successful genome editing was achieved for five of eight genes: CTRP, Pfs47, Pfs25, WARP, and HSP101. Attempts to edit SOAP, P230, and Pf3D7_1004600 using alternative guide RNAs were unsuccessful, as parasites did not recover post-transfection despite multiple efforts.

### Two genes show higher infection with sympatric parasite-allele and vector-species combinations

Based on our hypothesis that these genes are involved in regional adaptation to vector communities, we expected that replacement of the reference allele with the alternative allele should increase infectivity in species that are sympatric for the alternative allele and decrease infectivity in allopatric ones. To test this, we performed Standard Membrane Feeding Assays (SMFAs) using five allelic-replacement lines and their silent controls across four *Anopheles* species with distinct geographic ranges (**Figure 1A**). We defined a sympatric combination of parasite allele and vector species as a co-occurrence of the alternative allele at greater than 50% frequency in the broad geographic region (**Figure 1B**) and presence of the vector in at least one country within that region where parasite genomic information was available (**Figure 1A**, **S2**).

In total, 4,708 mosquitoes were infected across all experiments. Exflagellation rates prior to infectious feeding showed no differences between replacement and silent control lines (**Figure S4**), confirming comparable male gametocyte quality. Infection levels were assessed at the oocyst stage by measuring prevalence (proportion infected) and intensity (infection level among infected mosquitoes) (**Figure 2**). Oocysts were counted and measured 10 days post-infectious feed from dissected midguts. *An. gambiae* infected with the unmodified 3D7 clone was included as a positive control to confirm successful infection in each experimental replicate (**Figure S5**). We observed a reduction in infectivity in all edited lines compared to the 3D7 control, which may reflect effects associated with transfection, drug selection, an extended time in culture, and/or clonal selection rather than the specific genetic modifications (**Figure S5**).

**Figure 2.**
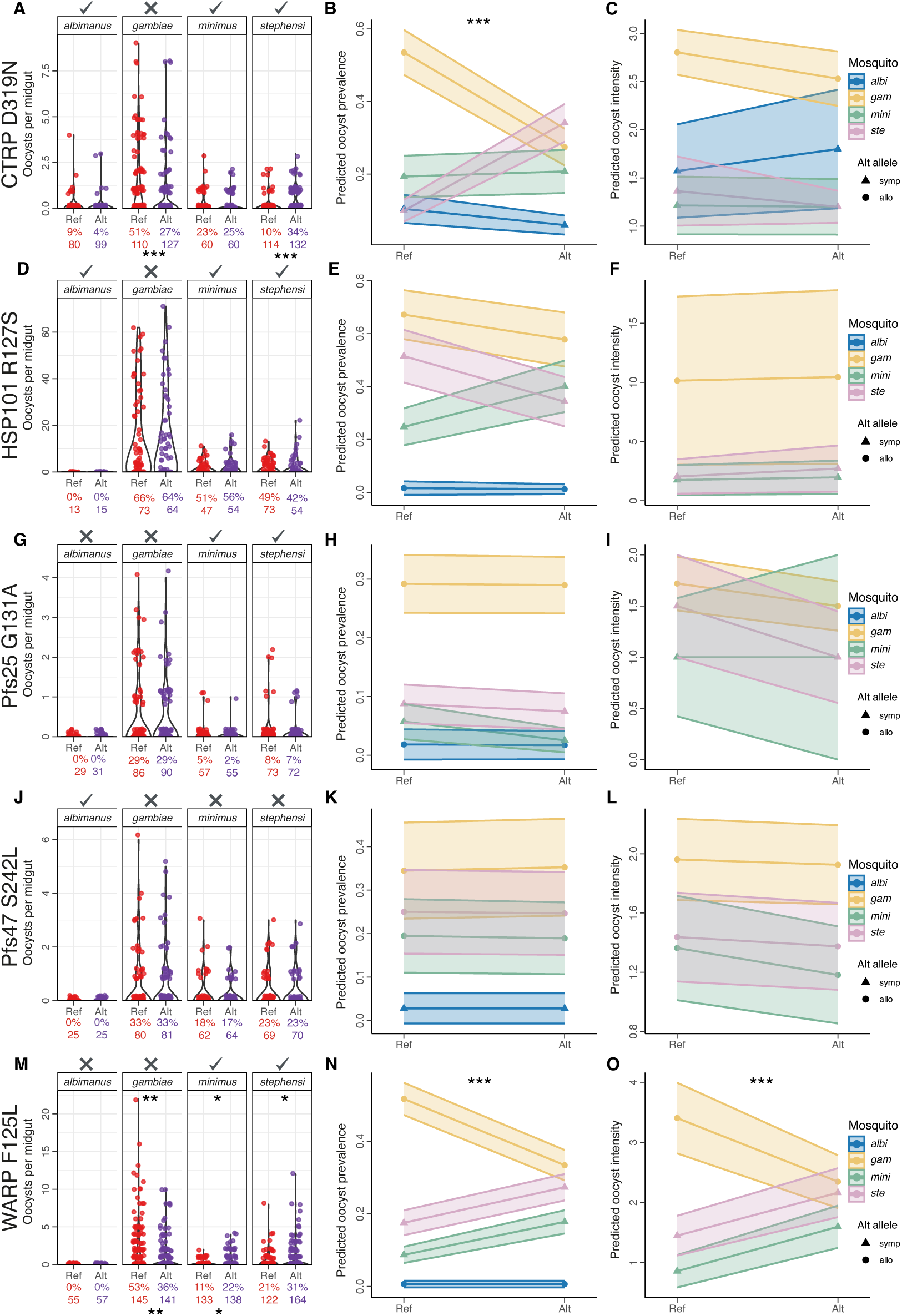
Infection of different *Anopheles* species by five *P. falciparum* allelic-replacement lines at oocyst stage. For each gene tested, the oocyst count data, predicted oocyst prevalence, and predicted oocyst intensity are displayed (CTRP: A-C; HSP101: D-F; Pfs25 G-I; Pfs47: J-K; WARP: M-O). The oocyst count data shows the number of oocysts for each mosquito midgut counted 10 days post infectious feed across experimental replicates. The violin plot shows all oocyst data for each species (*An. albimanus*, *An. gambiae*, *An. minimus*, *An. stephensi*) and each parasite allele (reference (Ref) vs alternative (Alt)). The proportion of infected mosquitoes and total number of midguts assessed is displayed below each violin. Stars represent significant differences between the reference and alternative allele within an individual mosquito species based on post-hoc pairwise comparisons from oocyst prevalence models (stars below) or oocyst intensity models (stars above). Sympatric combinations of alternative allele and vector species are are marked with a ✓ above the plot, whereas allopatric combinations are marked with an ✗ (A, D, G, J, M). The predicted prevalence plots show the estimated marginal means from the oocyst prevalence model (logistic regression) for each gene plotted by species and allele. Stars represent a significant Mosquito x Allele interaction for that locus (B, E, H, K, N). The predicted intensity plots show the estimated marginal means from the oocyst intensity model for each gene plotted by species and allele. Stars represent a significant Mosquito x Allele interaction (C, F, I, L, O). Points in predicted prevalence and intensity plots are shaped by whether the alternative allele is sympatric (triangle) or allopatric (circle) for that mosquito species. Shaded regions around each reaction norm represent the standard error. A minimum of three experimental replicates was performed for each gene. *** p <0.001, ** p <0.01 * <0.05.

Significant mosquito-by-allele interactions were detected for two genes, CTRP and WARP, with sympatric combinations showing higher infection levels compared to allopatric ones. Both genes showed a mosquito-by-allele effect on oocyst prevalence (*p* < 0.001; **Figure 2B**, **2N**). For CTRP D319N, oocyst prevalence was reduced in *An. gambiae* (allopatric; adj *p* < 0.001) but elevated in *An. stephensi* (sympatric; adj *p* < 0.001; **Figure 2A**). Similarly, the WARP F125L allele increased oocyst prevalence in *An. minimus* (sympatric; adj *p* = 0.018) and decreased it in *An. gambiae* (allopatric; adj *p* = 0.003; **Figure 2M**).

WARP also showed a significant mosquito-by-allele effect on oocyst intensity (*p* < 0.001; **Figure 2O**), with the WARP F125L allele yielding higher infection loads in *An. minimus* (sympatric, adj *p* = 0.047) and *An. stephensi* (sympatric, adj *p* = 0.040) and lower loads in *An. gambiae* (allopatric, adj *p* = 0.003) (**Figure 2M**). No significant differences in intensity were observed for CTRP D319N. The remaining alleles (HSP101 R127S, Pfs25 G131A, and Pfs47 S242L) had no measurable effects on oocyst prevalence or intensity (**Figure 2D-L**).

For CTRP and WARP, an additional clone of both the allelic-replacement and silent control was tested to confirm the vector-specific transmission phenotype. We observed the same pattern of the allele in each species as seen with the primary clones (Figure 3A). We also measured oocyst diameter in a subset of the infected midguts and found no mosquito-by-allele effect on oocyst size at day 10 post-infectious feed in CTRP or WARP (**Figure 3B**) or for the other three genes tested (**Figure S6**). We then quantified sporozoite prevalence and intensity in the mosquito salivary glands at day 17 post infection using qPCR for CTRP and WARP and found a significant mosquito-by-allele interaction for WARP (*p* = 0.034), with the alternative allele (F125L) showing lower sporozoite prevalence compared to the reference in *An. gambiae* (allopatric, adj *p* = 0.018, **Figure 3C**). Similarly, CTRP D319N was associated with reduced sporozoite prevalence in *An. gambiae* (adj *p* = 0.027, **Figure 3C**). No significant mosquito-by-allele interaction was found for sporozoite intensity for either gene. Surprisingly, sporozoite prevalences were generally higher than oocyst prevalences, and sporozoites were detected in *An. albimanus* infected with the WARP-F125 parasite, but oocysts were not. These differences may reflect variation in assay sensitivity or specificity, and recent work has seen the same discrepancy (higher sporozoite prevalence compared to oocysts) using the same qPCR primers, and suggests there is a higher false-positive rate in the qPCR assay [36]. Despite this discrepancy, we note the trend seen across the two genes in *An. albimanus*, where the alternative allele showed higher sporozoite prevalence in the sympatric combination compared to its reference (CTRP D319N-*An. albimanus*) but allopatric combination (WARP F125L- *An. albimanus*) resulted in a decrease in sporozoite prevalence (**Figure 3C**).

**Figure 3.**
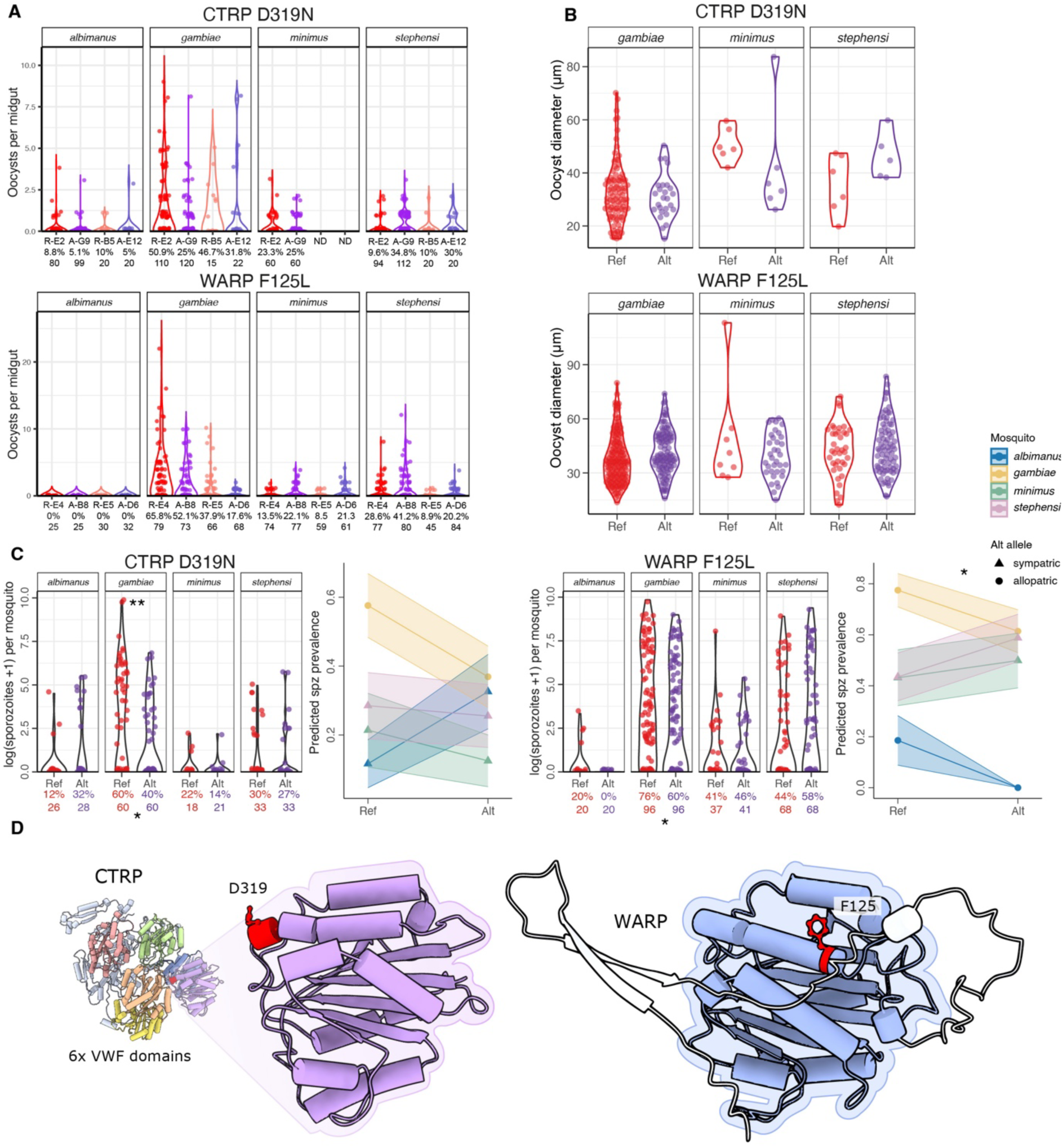
CTRP D319N and WARP F125L show vector-species dependent transmission levels and are in von Willebrand factor A (vWFA) domains. (A) Oocysts per midgut are shown by clone. For CTRP, Ref E2 and Alt G9 were the primary clones tested and Ref B5 and Alt E12 were the secondary clones tested. For WARP, Ref E4 and Alt B8 were the primary clones tested, whereas Ref E5 and Alt D6 were the secondary clones tested. Each dot represents oocyst counts from a single mosquito midgut. No significant allele-by-mosquito-by-clone interactions were observed for oocyst prevalence or intensity for either gene. (B) Oocyst diameter for CTRP and WARP alleles across vector species. No significant mosquito-by-allele interaction was observed. (C) Sporozoite infection levels for CTRP and WARP alleles. For each gene, the left panel shows the the violin plot shows all sporozoite data for each species (*An. albimanus*, *An. gambiae*, *An. minimus*, *An. stephensi*) and each parasite allele (reference (Ref) vs alternative (Alt)). The proportion of infected mosquitoes and total number of midguts assessed is displayed below each violin. Stars represent significant differences between the reference and alternative allele within an individual mosquito species based on post-hoc pairwise comparisons from sporozoite prevalence models (stars below) or sporozoite intensity models (stars above) ** p <0.01, * <0.05. The right panel shows the estimated marginal means from the sporozoite prevalence model for each gene is plotted by species and allele. A star represents a significant Mosquito x Allele interaction (p <0.05). Points are shaped by whether the alternative allele is sympatric (triangle) or allopatric (circle) for that mosquito species. Shaded regions around each reaction norm represent the standard error. (D) The protein structure of CTRP and WARP predicted by UCSF Chimera software [45]. Distinct vWFA domains are shown in colour. The D319N substitution in CTRP reduces negative charge on this residue (negative charge from aspartic acid replaced with uncharged asparagine). The F125L substitution in WARP removes an aromatic side chain (phenylalanine replaced by leucine).

### CTRP D319N and WARP F125L both fall in von Willebrand factor A domains

Protein structure analysis of WARP and CTRP revealed that CTRP contains six von Willebrand factor A (vWFA) domains, whereas WARP has a single vWFA domain (**Figure 3D**)[29,31,32,37]. Both CTRP D319N and WARP F125L substitutions occur within these domains. The vWFA domains of CTRP are essential for gliding motility and onward transmission through the mosquito [38]. By homology to integrin-like adhesion modules, these domains are predicted to mediate attachment to host surfaces through metal ion–dependent adhesion sites (MIDAS) [39,40].

The D319N substitution in CTRP replaces a negatively charged aspartic acid with an uncharged asparagine (**Figure 3D**). Aspartic acid residues within vWFA domains may coordinate bound cations at or near the MIDAS motif, which could stabilize ligand binding and domain conformation [41,42]. Loss of this negative charge may weaken cation coordination and alter the structural dynamics of the vWFA domain, potentially modifying adhesion to mosquito midgut epithelial ligands or affecting downstream processes required for ookinete motility and invasion [27,32,38].

Similarly, the WARP F125L substitution replaces phenylalanine with leucine, removing an aromatic side chain (**Figure 3D**). Aromatic residues often contribute to hydrophobic packing within vWFA cores and π-interactions that stabilize protein–protein interfaces [43,44]. The loss of this aromatic interaction may subtly affect domain folding or reduce affinity for midgut-associated receptors such as basal lamina components and mosquito-derived lectins [31].

Given that vWFA domains mediate critical adhesion processes, these substitutions are likely to modulate protein–protein or protein–matrix binding in the midgut. We hypothesize that such alterations could underlie the observed allele-specific differences in transmission efficiency by tuning parasite–vector compatibility at the molecular interface of midgut invasion.

## Discussion

Understanding how spatial variation and local adaptation shape *P. falciparum* genetic diversity and transmission dynamics is fundamental to public health because treatment or intervention in one location may be ineffective in another. Here we have demonstrated how natural allelic variation in *P. falciparum* transmission-stage genes can produce vector-specific effects on parasite infectivity. By combining population genomic data with functional assays across multiple geographically diverse *Anopheles* species, we show that two variants, CTRP D319N and WARP F125L, modulate infection outcomes in a manner consistent with adaptation to sympatric vector hosts. These results provide direct experimental support for the long-standing hypothesis that *P. falciparum* is adapting to regional vector community composition. Additionally, our results challenge the lock-and-key paradigm that compatibility is determined by a single *P. falciparum* locus (Pfs47), but instead indicate that this is a complex, polygenic trait where multiple molecular interactions determine vector specialisation.

CTRP and WARP encode micronemal proteins required for ookinete motility and midgut invasion and both contain vWFA domains that are likely involved in adhesion to the midgut epithelium and contained the polymorphisms of interest. The specific amino acid changes within the vWFA domains altered the charge of the residue for CTRP and removed an aromatic ring for WARP. These changes could potentially modify the strength or the specificity of binding to mosquito midgut ligands such as laminins, lectins or glycans [31,46,47]. Future work to identify the binding partners of *P. falciparum* WARP and CTRP, and how the strength and specificity of these interactions change with different parasite alleles and sympatric vs allopatric vector midgut binding partners, could help elucidate the underlying molecular mechanisms driving the mosquito-by-allele phenotypes we observed.

We did not observe mosquito-by-allele interactions for Pfs47, HSP101, or Pfs25. We cannot rule out that these loci are also regionally adapted to different vectors, given the limited number of mosquito species tested and the fact that only one parasite genetic background was tested. Additionally, for Pfs47, previous work has shown that the full haplotype with four amino acid changes is needed to match the regional vector (*An. gambiae* vs *An. albimanus*) [48]. However, the S242L mutation in an NF54 parasite background has previously been shown to result in immune detection and parasite melanisation by the *An. gambiae* L35 resistant strain [48]. This difference from our results is likely explained by the use of different *An. gambiae* strains, as we do not regularly observe melanisation in our *An. gambiae* Kisumu colony. For all loci tested, future investigations could employ a full haplotype replacement strategy, as we only edited a single amino acid substitution — the variant with the highest *Fst* that contributed to the global differentiation score [13]. Several additional non-synonymous SNPs showed similar fixation patterns within each gene (**Figure S7**), and together these may enhance compatibility between parasite and vector [35]. Additionally, future work could test multiple genetic backgrounds and combine haplotypes from different genes to understand additive and epistatic effects on vector compatibility. *P. falciparum* lineages from these distinct geographic areas are genomically very different, which is likely due to very different selective pressures beyond vector species including transmission intensity and drug-pressure [3,49,50]. Understanding how these selective pressures together shape the parasite genome and the spread of distinct lineages, including drug-resistant *P. falciparum*, is key to understanding the epidemiology of the disease.

Together, these findings demonstrate that naturally occurring amino acid substitutions in key transmission-stage proteins can alter *P. falciparum* infectivity in a vector-dependent manner. The effects of CTRP D319N and WARP F125L across mosquito species provide experimental evidence that parasite–vector compatibility is an evolving, polygenic trait shaped by spatially variable ecological interactions. Such adaptation has important implications for the geographic structuring of malaria transmission and for the design of transmission-blocking interventions. For example, as the distribution of *An. stephensi* expands into East Africa [51], its contribution to transmission could depend on the circulating parasite genotypes in this region. Accounting for natural allelic variation and vector diversity will be essential to developing control strategies that remain effective across the heterogeneous parasite and mosquito populations that sustain global malaria transmission.

## Materials and Methods

### Generation of Transgenic Parasites

The pBgC plasmids [52,53] were constructed by sequentially cloning the gRNA oligo and donor template. The plasmid was first digested with BsaI, then the sgRNA oligo was inserted by In-Fusion cloning (Takara Bio), following the manufacturer’s protocol. The construct was transformed into XL-10 Gold competent cells, and colonies containing the sgRNA insert were selected. A donor template (300–600 bp) containing the desired mutation along with silent mutations was synthesized and cloned into the HindIII and EcoRI sites of the plasmid using In-Fusion. After transformation, four colonies were selected for sequencing to confirm insertion of the correct donor template.

*P. falciparum* 3D7 parasites were transfected with pBgC plasmids containing the desired non-synonymous mutation or silent control for each gene (**Table S2**). Synchronized ring-stage parasites (5–10% parasitemia) were transfected with 80–100 μg of plasmid. Transfected parasites were selected with 2.5 μg/ml Blasticidin S for 8 days, then returned to normal 1640 medium supplemented with 5% human serum, 25 mM HEPES, 50 uM hypoxanthine and 0.2% sodium bicarbonate. Cultures were incubated at 37°C in a gas mixture of 1% O₂, 3% CO₂, and 96% N₂. Once parasites recrudesced, genomic DNA was extracted, amplified using the primers in **Table S3** and sequenced to confirm genome editing. Limiting dilution cloning was done to select clones that had the desired mutation. Bulk cultures were diluted to 0.5% prior to serial dilution in 96-well plate. After two weeks, 50 μl from each well was stained with 1:5000 SYBR Green in PBS to detect parasites by flow cytometery (MACSQuant). Clonal lines were expanded and sequenced to confirm successful editing.

### Gametocyte Induction

Gametocytes were induced at 0.5–0.7% asexual parasitaemia and 6% hematocrit in 25 cm^2^ flasks with 5 ml of culture following established protocols [54]. Two flasks were set up for the day 17 gametocytes and three days later two flasks were set up for the day 14 gametocytes. Flasks were incubated at 37°C and media was changed daily. The media was increased to 1.5x when parasitaemia was 3-5%.

### Mosquito rearing

*An. albimanus* PANAMA, *An. gambiae* KISUMU, *An. minimus* MINIMUS1, and *An. stephensi* SDA500 were reared for transmission experiments at 27 degrees C, 70% relative humidity and 12 hour by 12 hour light/dark cycle conditions. Larvae were fed on ground Tetramin and Tetra Pond pellets. Adults were maintained on a 5% D-glucose solution with 0.05% para-amino benzoic acid (PABA) via cotton pads placed on top of cages. *An. albimanus* were maintained on 10% glucose solution with 0.05% PABA.

### Standard Membrane Feeding Assay and Exflagellation

The Standard Membrane Feeding Assay (SMFA) [55,56] was performed by pooling gametocyte cultures from day 14 and day 17 flasks. The cultures were centrifuged at 500 × g for 7 minutes at 37 °C, then resuspended with an equal volume of human serum and diluted to 1% gametocytemia with 40% hematocrit. Blood meals were loaded in glass membrane feeders covered with goldbeater’s skin, connected to a 37°C water bath. Mosquito pots that had 4-7 day old female mosquitoes were put under the glass feeder for 10 minutes to feed and then maintained at 27°C with 70% humidity.

Exflagellation was assessed from an aliquot of the prepared blood meal by diluting to 2.5% haematocrit in ookinete medium containing xanthurenic acid (10.4 g RPMI powder, 2 g of NaHCO_3_, 50 mg of hypoxanthine in 1 L of dH_2_O and 1 ml of 100 mM xanthurenic acid) [57]. 10 ul of the dilution was added to a hemocytometer (Neubauer Improved C-Chip). After 10 minutes, exflagellation events were counted under 40X magnification for 10 minutes to estimate the number of events per μl at 1% gametocytemia.

### Oocyst and Sporozoite Quantification

Ten days post feeding, mosquitoes were killed with 70% ethanol. Midguts were dissected in PBS and stained with 1% mercurochrome (25 μl per sample). Midguts were examined at 20X magnification on an Echo revolve microscope. The number oocysts per midgut was counted and oocyst size was measured.

Mosquitoes were frozen at day 17 post infectious feed at -20 °C for sporozoite measurement using qPCR. DNA extraction was performed on dissected mosquito heads and top of thorax using the DNeasy Blood & Tissue Kit (Qiagen). Quantitative PCR was performed by mixing 1X SYBR Green Master Mix (Applied Biosystems) and 600 nM of each primer targeting the *PfCOX* gene (forward: 5’-TTACATCAGGAATGTTATTGC-3’, reverse: 5’-ATATTGGATCTCCTGCAAAT-3’, total reaction 15 ul) [58]. Quantitative PCR was run on a Viia 7 system with the following cycling conditions: denaturation: 98°C and annealing: 56°C. Melting curve analysis was used to confirm product specificity. Standard curves were generated using known numbers of ring-stage parasites; 100000, 10000, 1000, 100, 10 and 1.

### Statistical Analysis

Statistical models for oocyst and sporozoite prevalence and intensity, as well as oocyst size, had the following core structure for each gene:

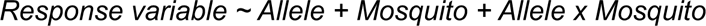

Where Allele represents the reference or alternative allele for that locus and Mosquito represents the vector species in which the oocysts or sporozoites were measured (*An. albimanus*, *An. gambiae*, *An. minimus*, or *An. stephensi*). For oocyst prevalence, logistic regression (family = binomial(link = “logit”)) was fitted with a biased-reduced generalized linear model using the brglm2 package in R with Replicate as a fixed effect in the model [59]. This allowed for analysis of mosquito-allele combinations that had no oocysts detected [60]. For oocyst intensity, non-zero samples were analysed using a generalized linear mixed model with a negative binomial distribution (family = nbinom2) using the glmmTMB package in R [61]. Replicate and Clone were considered as random effects for CTRP and WARP where two clones were tested, and Replicate alone was considered a random effect for HSP101, Pfs25, and Pfs47. For sporozoite prevalence, glmmTMB was used for logistic regression again with Replicate and Clone as random effects. For sporozoite intensity, glmmTMB was used to fit a generalised linear mixed model with a Gaussian distribution with the same random factors as the sporozoite prevalence model. For oocyst size models, a linear mixed effects model was fitted using the lme4 package in R with Midgut from which the oocyst was measured as a random effect [62]. Replicate was considered a fixed factor for CTRP, Pfs25, Pfs47 and WARP, but not for HSP101 as oocyst size was only measured for one of the three experimental replicates. To test the effect of clone, Clone and Clone x Allele x Mosquito were considered as fixed effects in the model. A reduced model was tested without with the Clone x Allele x Mosquito interaction and the full and reduced models were compared with an LRT test. For oocysts and sporozoite models, estimated marginal means and pairwise contrasts within species were calculated using the emmeans package in R [63].

### Vector distribution mapping

Rasters downloaded from the Malaria Atlas Project [2,34] are continuous probability of occurrence surfaces with values from 0 to 1. To produce a clear distribution footprint, surfaces were converted to a presence polygon using a fixed threshold of 0.5. Shaded areas indicate the probability of presence is higher than 50%.

### Protein structure visualisation

Protein structures were obtained from the AlphaFold database: AF-Q8IAL0-F1 (WARP) and AF-O97267-F1 (CTRP). Domain visualization, amino acid charge analysis, and mutation modelling were performed using UCSF ChimeraX version 1.7.

## Acknowledgements

We thank Luca Nelli for creation of the map in Figure 1A and Victor Tobiasson for support with the protein structure diagram in Figure 3D. We thank Farzana Khaliq, Andy Waters, and Katarzyna Modrzynska for production of the *An. stephensi* mosquitoes. We thank Francesco Baldini, Arthur Talman, and Adam Dobson for helpful feedback on earlier versions of this manuscript. This work was funded by a Sir Henry Dale Fellowship jointly funded by the Wellcome Trust and the Royal Society (Grant 220185/Z/20/Z) and a Lord Kelvin Adam Smith Leadership Fellowship awarded to VMH.

## Supplementary Information

### Supplementary Figures

**Figure S1.**
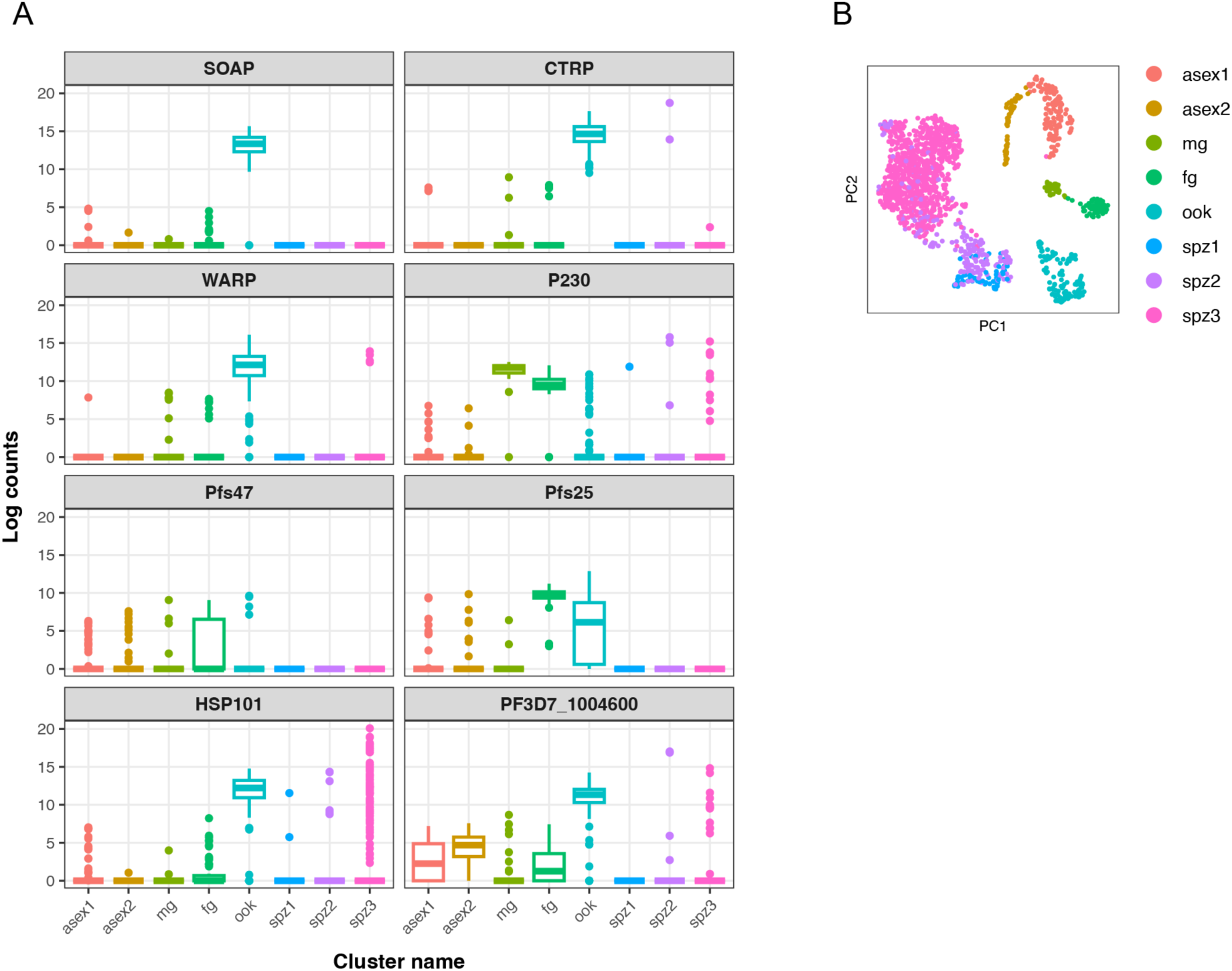
Expression of candidate genes in Malaria Cell Atlas data. Expression of the eight candidate genes across Malaria Cell Altas Smart-Seq2 *Plasmodium falciparum* data from [15,16]. Expression is shown for each gene in each stage (A) based on annotated clusters (B). Candidate genes showed high expression in the mature gametocytes or ookinete stages. Annotated clusters are ordered by developmental progression. Asex = asexual, mg = male gametocyte, fg = female gametocyte, ook=ookinete, and spz = sporozoite. Targeted candidate genes were selected from the intersection of genes in the top 5% of global differentiation scores from MalariaGEN data [13] and genes in the Malaria Cell Atlas data that either fell in gene clusters 7-12 or were more highly expressed in *P. falciparum* ookinetes compared to *P. berghei* (Table S1). Gene clusters 7-12 (g7-g12) are associated with high expression in *P. falciparum* gametocytes or ookinetes [15].

**Figure S2.**
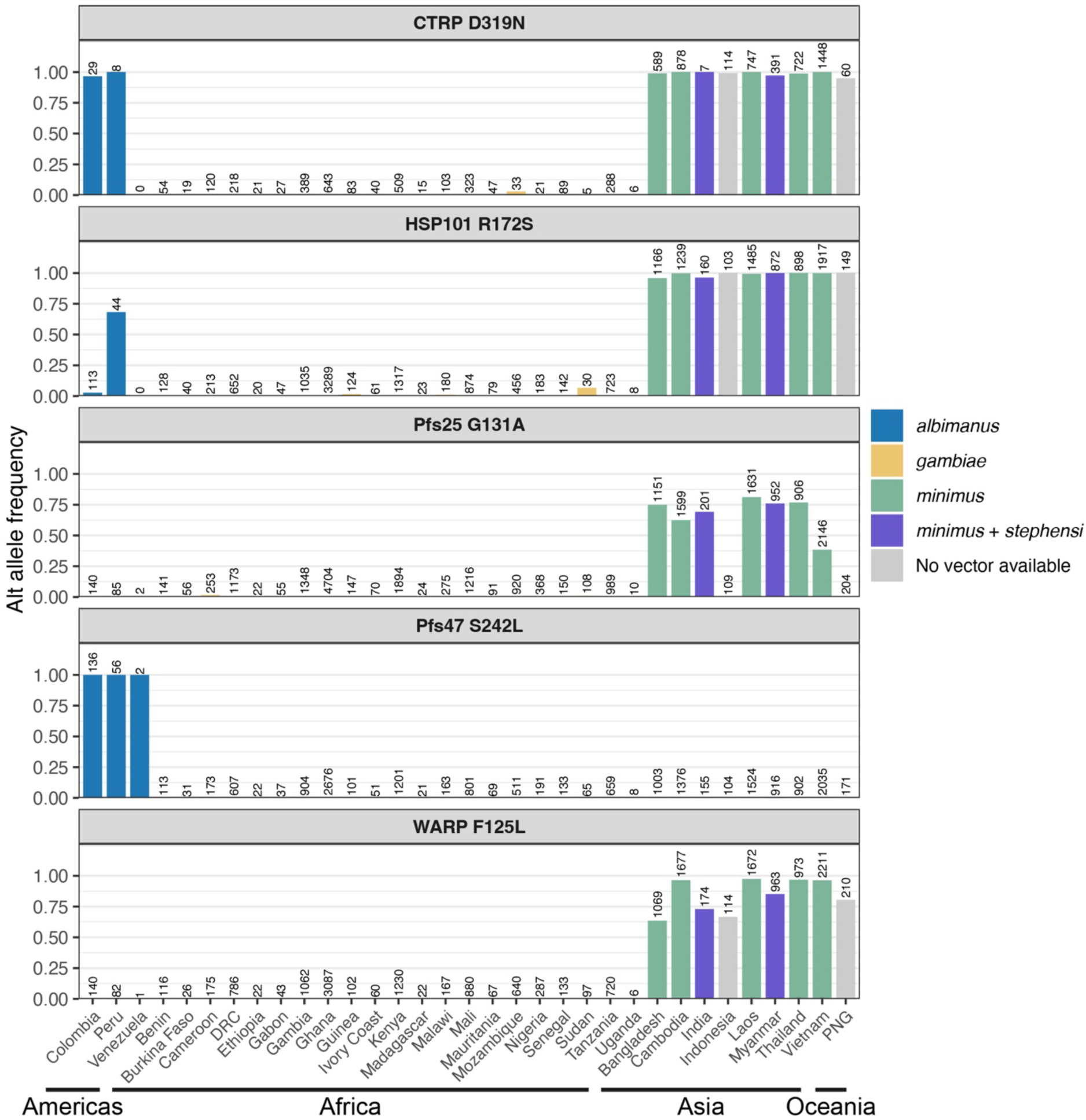
Alternative allele frequencies for each locus by country. Data are from MalariaGEN Pf8 [35] downloaded from HaploAtlas [64]. Bars are coloured by the vector or vectors that are present within that country according to Malaria Atlas Project data [2] and available as an established colony in our insectary. The number above each bar represents the number of samples from that country that passed QC and were maintained in the analysis set and HaploAtlas dataset for this gene by MalariaGEN [64].

**Figure S3.**
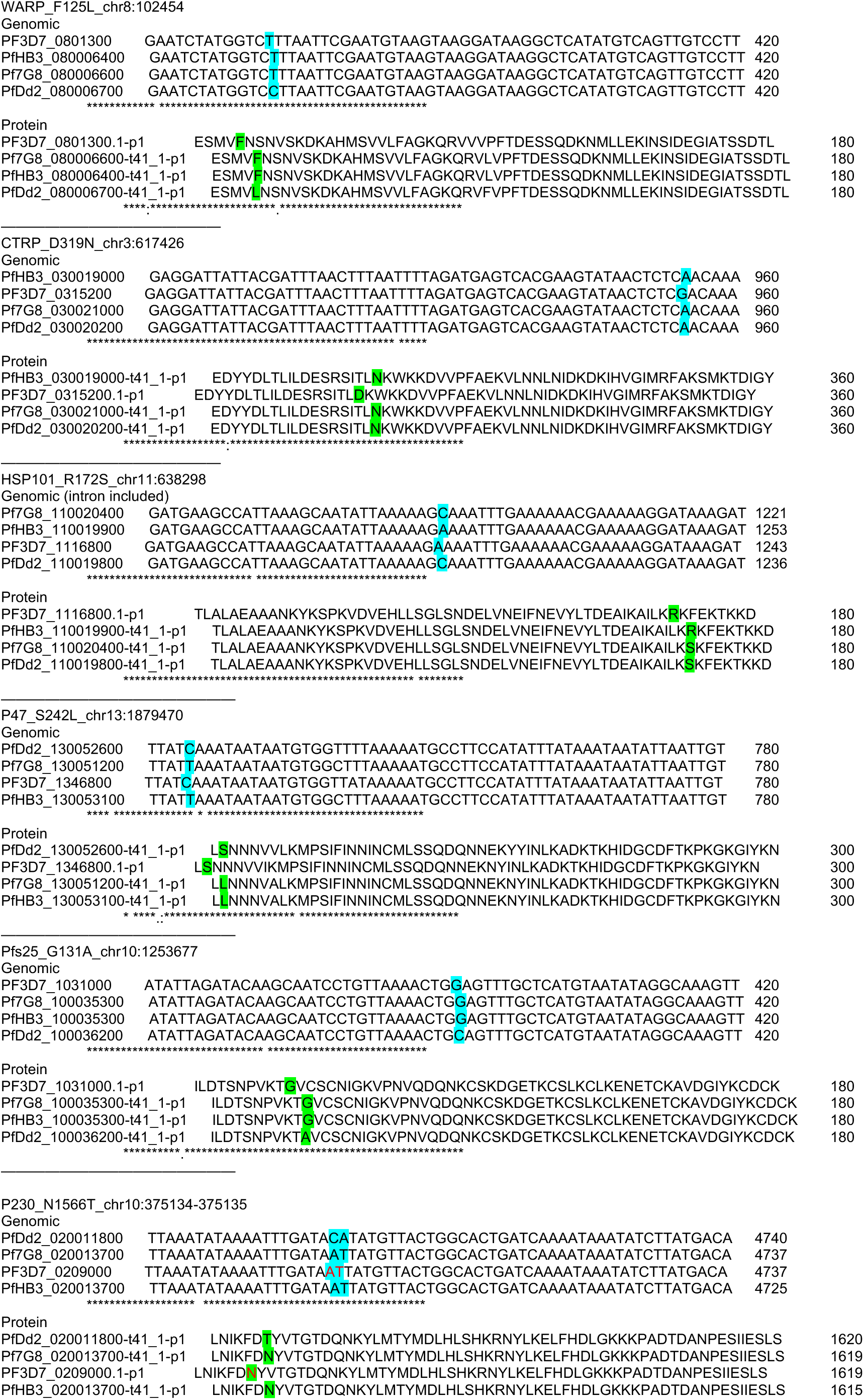
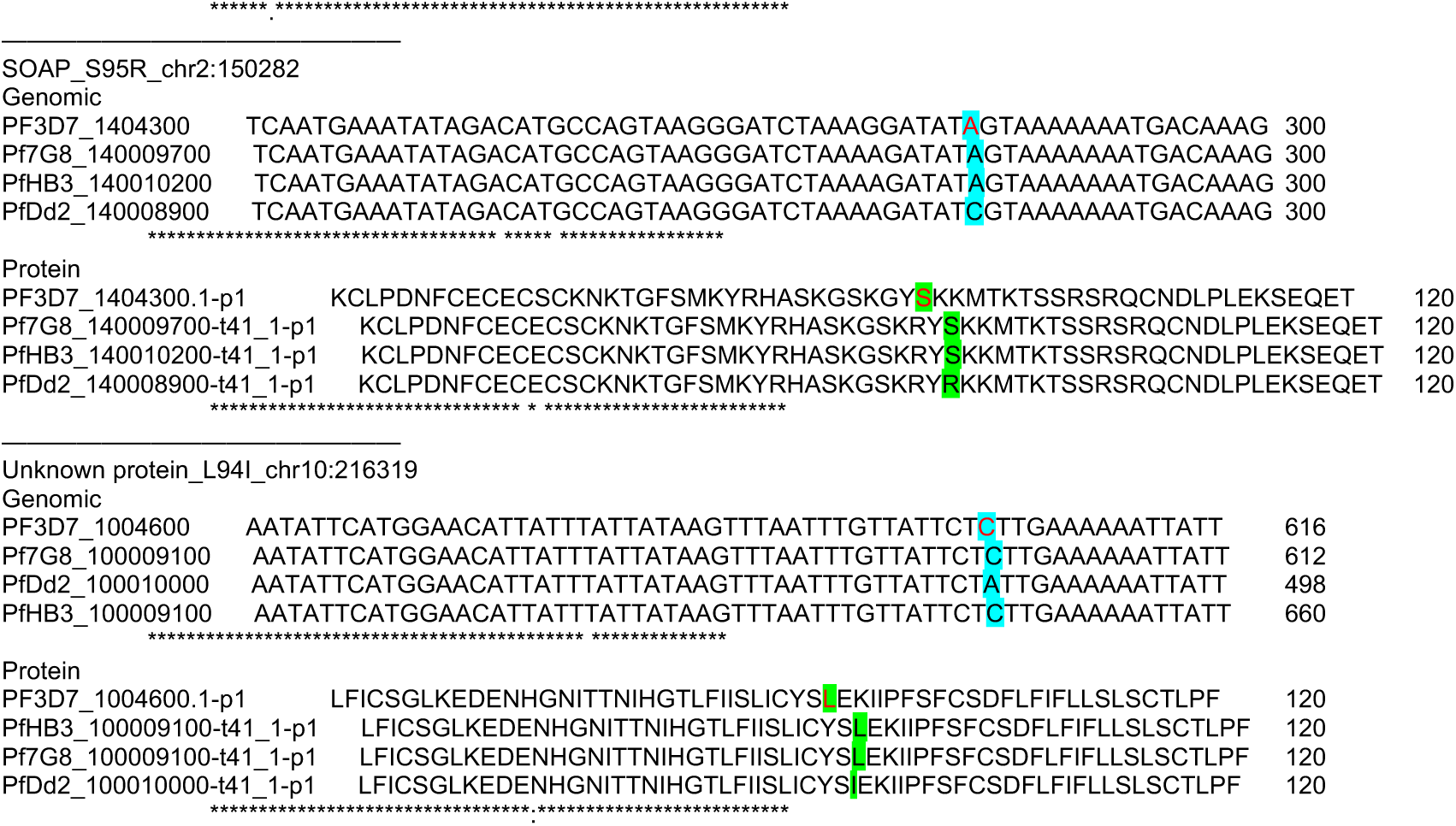
Genome and protein alignments for candidate genes. Non-synonymous SNPs identified in each gene selected for this study in four *P. falciparum* genomes from Africa (PF3D7), America (PfHB3 and Pf 7G8) and Asia (PfDd2). Targeted amino acids are highlighted in green within the protein sequence alignment, while blue indicates the genomic variants.

**Figure S4.**
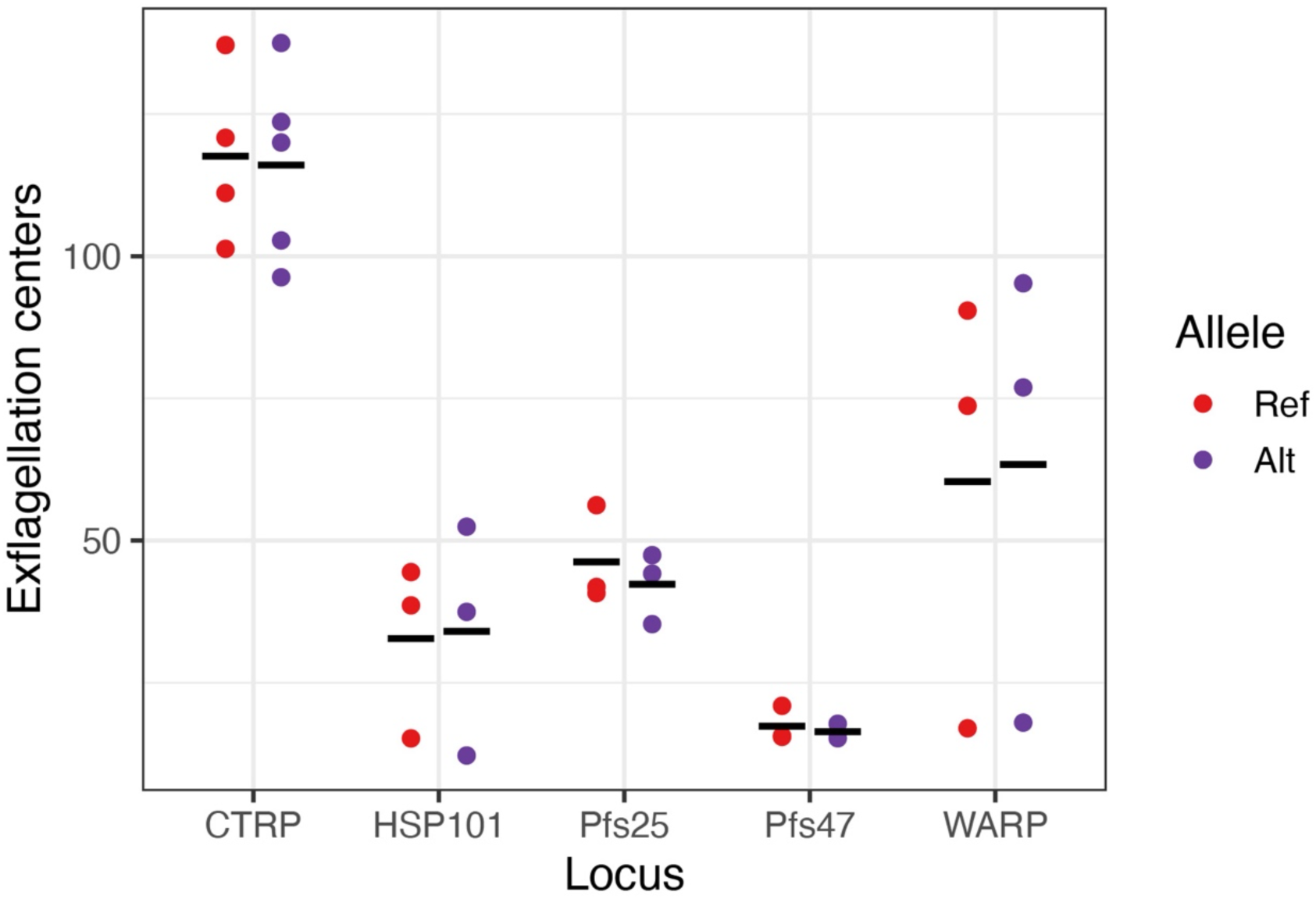
Exflagellation from five allelic-replacement parasite lines and silent controls. Exflagellation was measured for each infectious blood meal. Each point represents a single replicate, and the black bar represents the mean. No difference was observed between reference and alternative alleles in any of the five candidate loci (Linear model: Exflagellation centers ∼ Allele, *p* > 0.05 for all loci).

**Figure S5.**
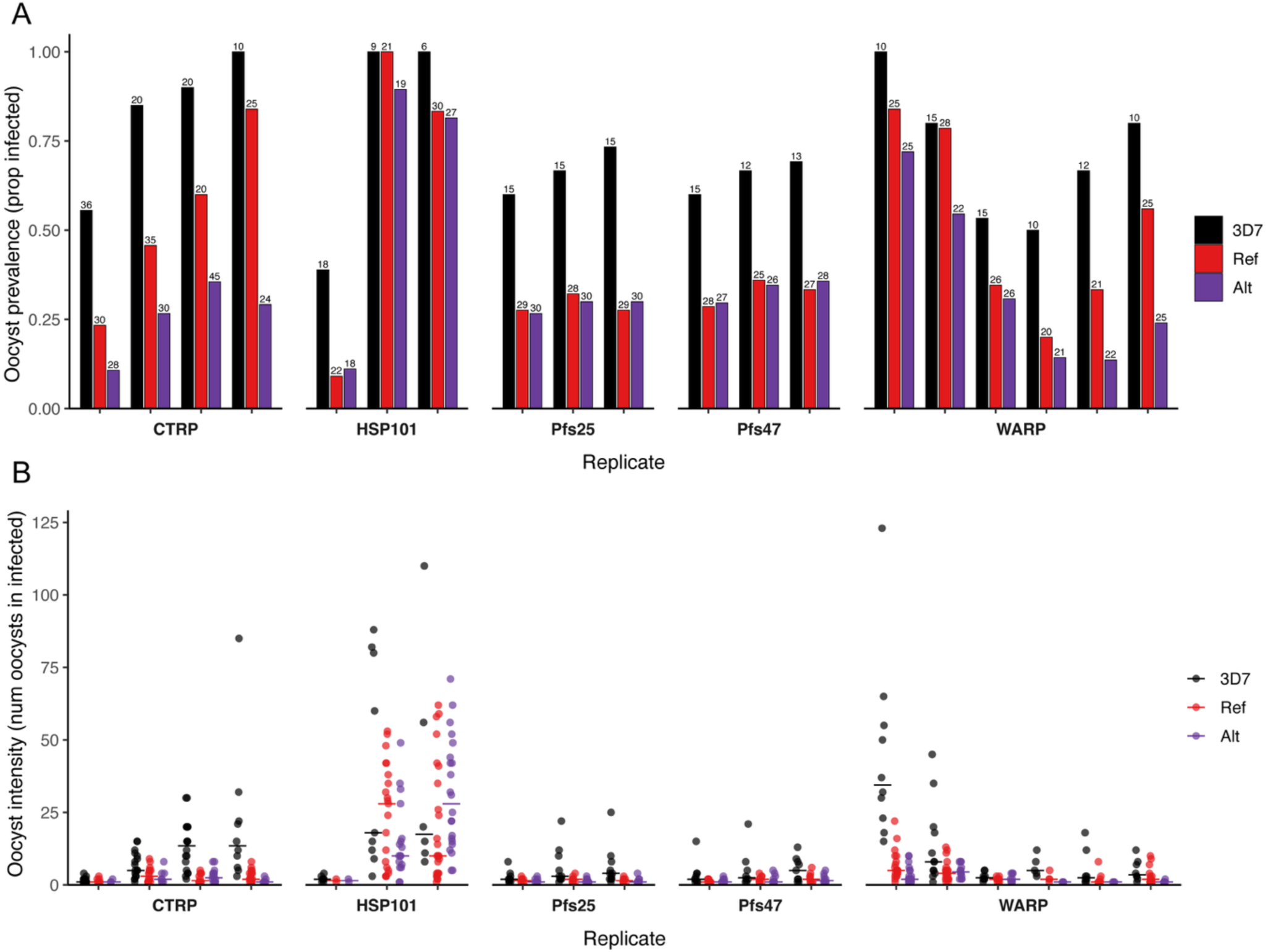
Oocyst prevalence and intensity for wild-type 3D7 and modified Ref and Alt lines for each gene in *An. gambiae* in each experimental replicate. *An. gambiae* infected with unedited wild-type 3D7 was included as a control in all experimental replicates to confirm that infection was achieved. (A) Shows the proportion infected in 3D7 alongside the edited reference and alternative line for each gene in each experimental replicate. The number above each bar represents the number of midguts examined. (B) The intensity of 3D7, Ref and Alt for each gene in each replicate. The bar represents the median.

**Figure S6.**
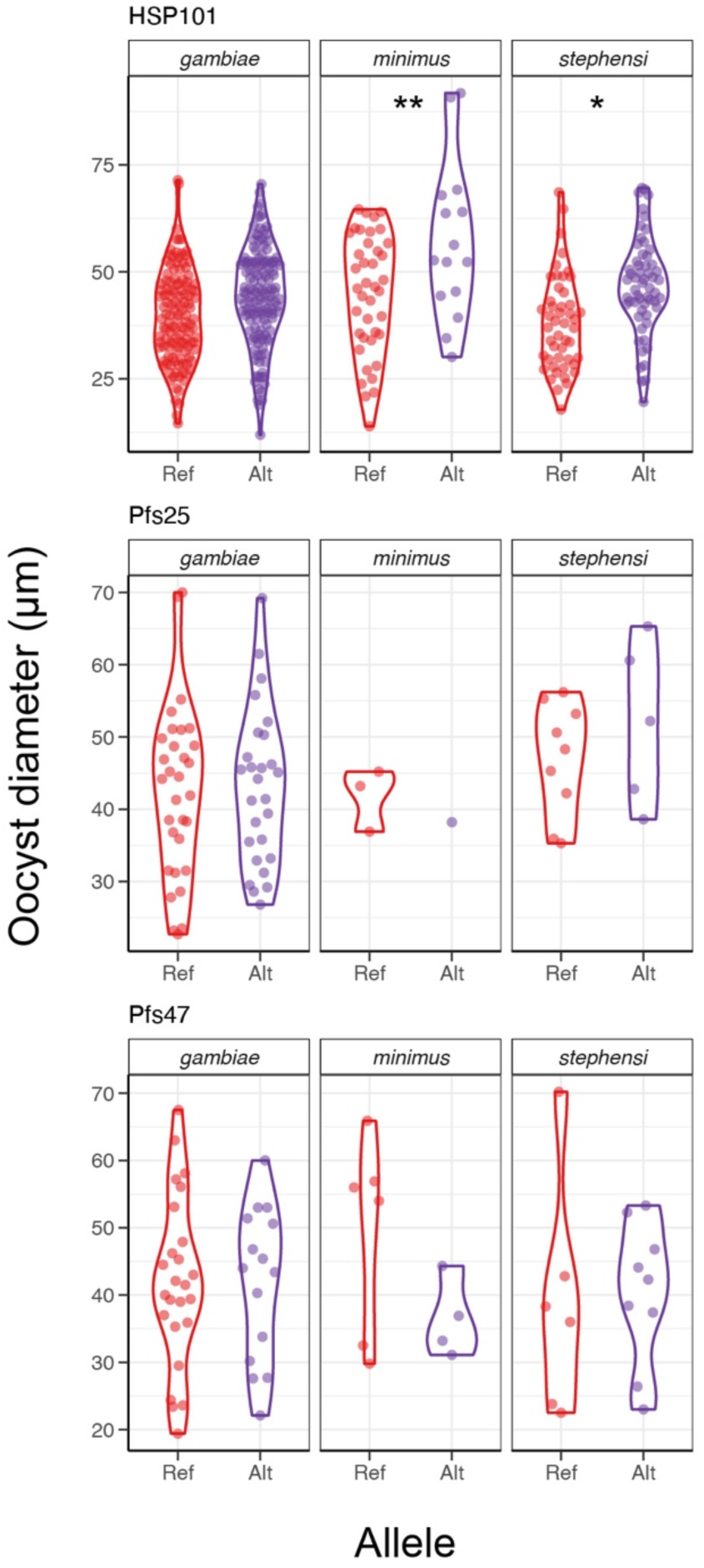
Oocyst diameter across parasite lines and vector species. No significant mosquito-by-allele interaction was found for any locus. However, for HSP101, the alternative allele had larger oocysts for *An. minimus (*adj *p* = 0.003) and *An. stephensi* (adj *p* = 0.040). Only a single replicate was measured for HSP101 compared to two or three replicates for all other loci.

**Figure S7.**
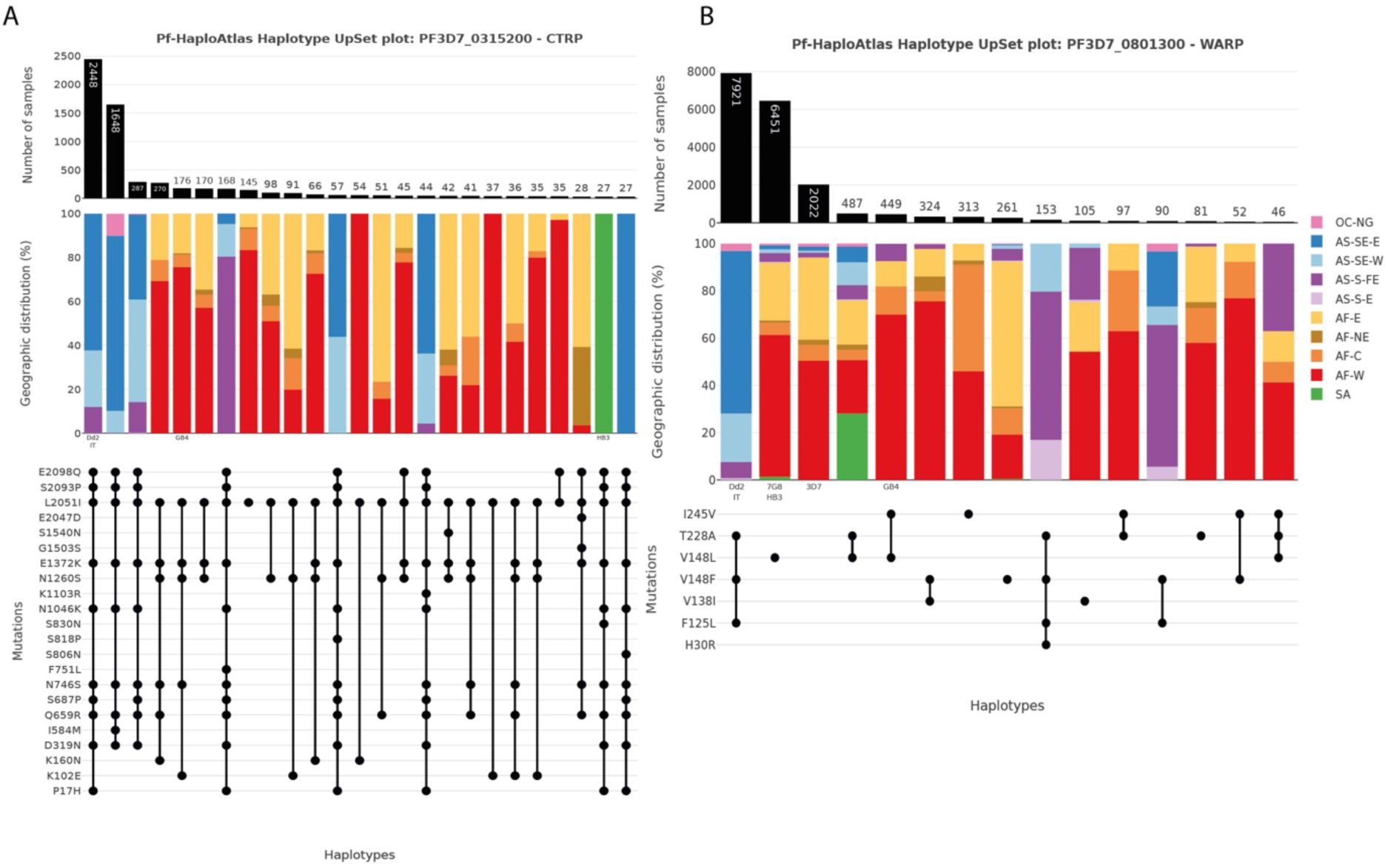
CTRP and WARP haplotypes. Images were created using the MalariaGEN Pf-HaploAtlas [64]. See https://pf-haploatlas.streamlit.app/?gene_id=PF3D7_0315200 and https://pf-haploatlas.streamlit.app/?gene_id=PF3D7_0801300 for more information. Legend terms: OC-NG = Oceania, New Guinea; AS-SE-E = Southeast Asia (East); AS-SE-W (Southeast Asia (West)); AS-S-FE = South Asia (Far East); AS-S-E = South Asia (East); AF-E = Africa (East); AF-NE = Africa (North-East); AF-C = Africa (Central); AF-W = Africa (West); SA = South America.

### Supplementary Tables

Table S1. 60 Candidate genes

See sup file

Table S2. Donor template sequences for mutant and silent control plasmids

See sup file

**Table S3.**
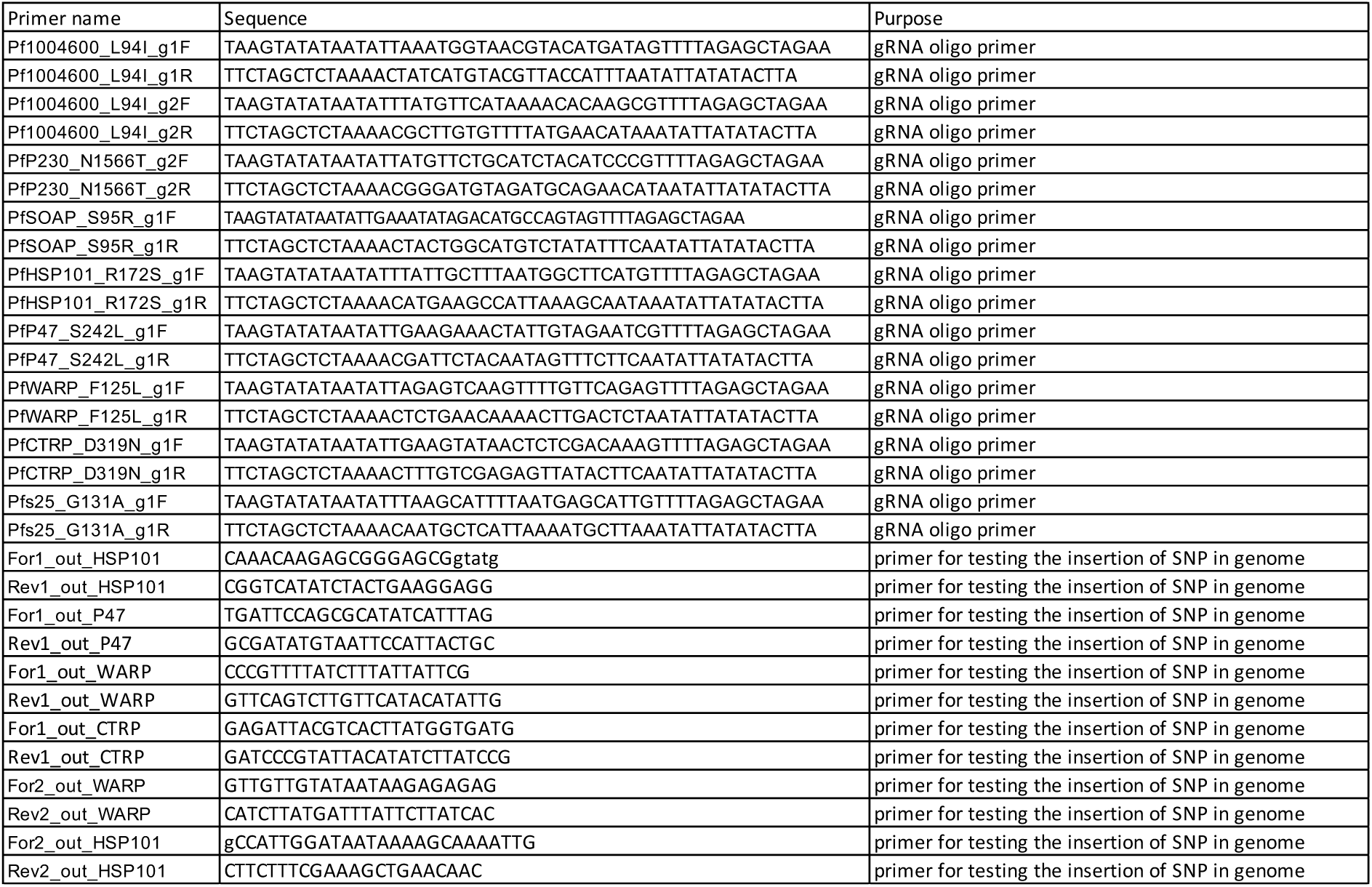
Guide oligonucleotide and primer sequences. The guide oligos were cloned into a plasmid construct to facilitate targeted gene editing, and the primer sets were employed to confirm the successful insertion of the SNP.

